# Low dimensional criticality embedded in high dimensional awake brain dynamics

**DOI:** 10.1101/2023.01.05.522896

**Authors:** Antonio J. Fontenele, J. Samuel Sooter, V. Kindler Norman, Shree Hari Gautam, Woodrow L. Shew

**Affiliations:** UA Integrative Systems Neuroscience Group, Department of Physics, University of Arkansas, Fayetteville, AR, USA, 72701

## Abstract

Whether cortical neurons operate in a strongly or weakly correlated dynamical regime determines fundamental information processing capabilities and has fueled decades of debate. Here we offer a resolution of this debate; we show that two important dynamical regimes, typically considered incompatible, can coexist in the same local cortical circuit by separating them into two different subspaces. In awake mouse motor cortex, we find a low-dimensional subspace with large fluctuations consistent with criticality – a dynamical regime with moderate correlations and multi-scale information capacity and transmission. Orthogonal to this critical subspace, we find a high-dimensional subspace containing a desynchronized dynamical regime, which may optimize input discrimination. The critical subspace is apparent only at long timescales, which explains discrepancies among some previous studies. Using a computational model, we show that the emergence of a low-dimensional critical subspace at large timescale agrees with established theory of critical dynamics. Our results suggest that cortex leverages its high dimensionality to multiplex dynamical regimes across different subspaces.

**Teaser:** Temporal coarse-graining reveals a low-dimensional critical subspace coexistent with a desynchronized subspace in awake cortex.

## Introduction

The ongoing dynamics of neuronal populations in cerebral cortex are high dimensional and the nature of the dynamics differs across dimensions. Some dimensions contain large amplitude fluctuations, while other dimensions tend to have smaller fluctuations. Accumulating evidence shows that a relatively small fraction of the total possible dimensionality - i.e. a low-dimensional subspace - contains much of the large amplitude collective fluctuations (*1–6*). This fact is commonly used to justify simplified views of population activity using just a few dominant dimensions defined by principal component analysis (PCA), for example (*7, 8*). Less commonly discussed are the smaller amplitude, more desynchronized fluctuations in the higher dimensional subspace, beyond the first few principal components. These basic observations lead to an interesting possible resolution of a long-standing debate in systems neuroscience.

The debate concerns the answer to a fundamental question: what is the dynamical regime of cortical neural networks in awake animals? Some researchers argue in favor of a desynchronized dynamical regime with weak collective fluctuations. This view is supported by measurements of low correlations of spikes recorded from pairs of neurons. Also, the desynchronized regime comes with potential computational benefits, like low noise and high input discrimination (*9–11*). Others argue for the importance of synchronized dynamics, based on observed oscillatory brain activity and potentially improved long range signal transmission (*12–14*). Of particular importance for our work here, a third camp argues in favor of criticality - a dynamical regime poised at the boundary between uncorrelated and strongly coordinated regimes (*15–19*). Criticality manifests with moderate correlations, highly diverse spatiotemporal fluctuations, and is thought to confer multiple functional advantages, like optimal information transmission and dynamic range (*20–26*).

How might consideration of the high-dimensional geometry of population dynamics help resolve the debate about cortical dynamical regime? Here we hypothesize that cortical populations could support more than one dynamical regime at the same time by separating different regimes into different subspaces. Criticality could occupy the low-dimensional subspace with large fluctuations (first few principal components) while a more desynchronized regime could occupy the higher-dimensional subspace with weak fluctuations (principal components beyond the first few). If this possibility is correct, then a cortical circuit need not choose between weak and strong correlations; it can have both simultaneously.

Is this hypothesis consistent with previous studies? In particular, how could this hypothesis be consistent with the multiple previous studies that have analyzed recordings of spikes from awake cortex and concluded that the activity was not consistent with criticality (*27–32*). These studies found activity that appeared to be weakly correlated, more consistent with a desynchronized regime. At first glance, these studies seem to contradict our hypothesis here of a low-dimensional critical subspace with large, correlated fluctuations; why might these studies miss the critical subspace? A likely answer to this question comes from a comprehensive comparison of all attempts to seek evidence for criticality based on recordings with single neuron resolution in awake animals (22 cases in all including the 13 negative cases in the papers cited above, a complete list to our knowledge at the time of writing this paper, Supplementary Table S1). A distinguishing feature of the 9 cases that reported positive evidence for criticality was that they focused on relatively long time scale fluctuations; they considered coarse-grained spike counts in time bins larger than about 10 ms. This temporal coarse-graining was either a deliberate choice in the data analysis (*31, 33*) or was due to limited time resolution of experimental measurements, which is typical for calcium imaging (*32, 34–38*). The most strongly negative reports were based on analyses at the millisecond time scale, with little temporal coarse-graining. Based on this small meta-analysis of previous work and theoretical support for the importance of temporal coarse-graining (more on this in the results below), we refine our hypothesis. We hypothesize that desynchronized and critical subspaces coexist, but that the critical subspace is not apparent without temporal coarse-graining. To test this hypothesis we recorded spike activity from motor cortex of awake mice and compared our measurements to a ground-truth theoretical model of critical dynamics.

## Results

We performed extracellular spike recordings of up to 247 units in motor cortex of awake, behaving mice (Fig 1A, 4 mice, 19 recordings, n=104±43 single units, 44±18 multi-units per recording, 44±9 minutes recording duration, more details in Methods). Our analysis of each recording begins with generating an N × M spike count matrix (Fig 1B), where N is the number of neurons and M is the number of time bins (M = recording duration divided by time bin duration ΔT). The entry in the ith row and jth column is the number of spikes fired by the ith unit during the jth time bin. We performed principal component analysis on each spike count matrix, and found that the activity is high dimensional, but much less than N dimensional; 45±0.05% of principal components (PCs) were needed to explain 95% of variance (Fig 1C).

**Fig. 1.**
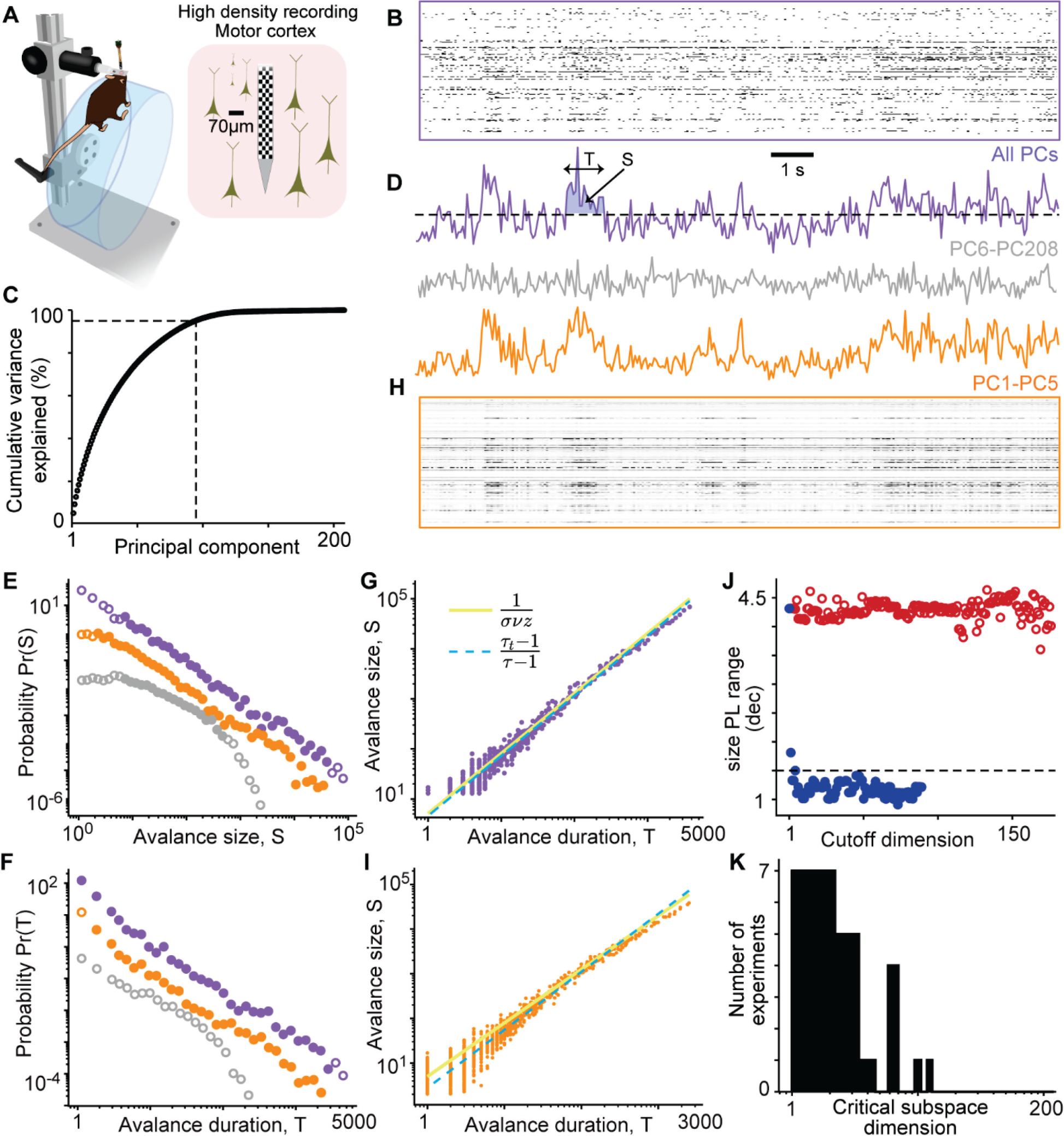
Low dimensional critical subspace. **(A)** Mice were head-fixed and placed on top of a wheel, free to run, rest, groom, etc. High density recordings were performed in motor cortex, using a Neuropixels 1.0 probe. **(B)** Example spike count raster plot for 208 units. **(C)** Cumulative variance of original activity explained by increasing number of principal components. Dashed lines marks the number of PCs needed to explain 95% of the variance. **(D)** Spike count time series for entire population (ΔT = 50 ms). Dashed line represents the avalanche threshold. The shaded suprathreshold excursion represents one avalanche. **(E)** Distributions of avalanche size (S) for the original data (purple), reconstructed data using the first five principal components (orange) and after removing the first five components (grey). Distributions are shifted vertically for comparison. **(F)** Same as in E, but for avalanche durations (T). **(G)** Avalanche sizes and durations follow predicted scaling law for original data. **(H)** Raster and population sum for reconstructed data based on the first five principal components (orange) or excluding the first five principal components (gray). **(I)** Scaling laws obeyed for PC1-5 reconstructed data. **(J)** Power-law range for avalanche size distributions as a function of PCs removed ascending/descending order (blue/red). Dimension of critical subspace is defined as the number of PCs removed before power-law range drops below 1.5 decades (dashed line). **(K)** Histogram of critical subspace dimension for all 19 recording sessions.

Next, we performed avalanche analysis following previously established methods (*26, 33, 39, 40*). We first created a 1-dimensional time series by summing spike counts across all N neurons. Then, we defined an avalanche as a period when this summed population activity exceeds a threshold (Fig 1D, see Methods). According to theory (*40, 41*), if the neural system operates near criticality, then avalanches should be diverse, with sizes (S) and durations (T) distributed according to power-laws, P(S) ∼ S^-τ^ and P(T) ∼ T^-τt^. We found good agreement with this prediction; avalanche sizes and durations were power-law distributed over a wide range of scales (Fig. 1E,F, purple). In addition, theory predicts that the exponents, τ and τ_t_, are not independent; they should be related to each other and to a third exponent (1/συz) according to the scaling law 1/συz = (1 - τ_t_)/(1-τ) (*40, 41*), sometime called the “crackling noise” scaling law. Moreover, according to theory, avalanche size depends on duration as S ∼ T^1/συz^. The avalanches we measured conformed well to all these predictions (Fig 1G). Thus, we conclude that the awake spike activity we observed here is in good agreement with predictions for a system operating at criticality. (As alluded to in the Introduction, this finding depends on the choice of time scale for the time bins ΔT, which we will investigate further below.)

## Critical subspace is spanned by first few principal components

Traditional avalanche analysis (Fig 1D-G purple) is based on the 1-dimensional population summed activity, which, by definition, precludes insight on how the dynamics may be distributed across the multiple dimensions revealed by PCA (Fig 1C). Do the scale-free avalanche dynamics exist in a subspace of the high dimensional population dynamics? If so, then the results of avalanche analysis should be robust to removal of PCs that are not in the avalanche subspace. We tested this possibility first with an example case (Fig 1 orange). We generated a new N × M activity matrix, reconstructed using only the first five PCs, excluding the other 203 dimensions (Methods). The population sum time series based on the reconstructed data was very similar to the original (Fig 1D,H).

Moreover, the avalanche statistics for this 5-dimensional reconstruction remained in good agreement with predictions for criticality (Fig 1 E,F,I orange). In contrast, if we reconstructed the data using PCs 6-208, the avalanche sizes and durations were not power-law distributed (Fig 1E,F gray). Thus, for this example, we conclude that the critical dynamics are primarily contained within the subspace defined by the first 5 PCs.

This example raises interesting questions. Is the dimensionality of this “critical subspace” exactly 5; is it higher or lower? Is the critical subspace always spanned by the first several PCs? To address these questions, we repeated the avalanche analysis using dimensionally-reduced reconstructed data, but systematically varying the cutoff dimension d_c_ from 1 to N (d_c_ = 5 in the example above). For each cutoff dimension we performed avalanche analysis on two data sets; one reconstructed using PCs 1 through d_c_, the other reconstructed using PCs d_c_+1 through N. For each case, we quantified the range of the avalanche size distribution that was well-fit by a power law. When reconstructed using PCs 1 through d_c_, the power-law range remained high, largely independent of d_c_ (Fig 1J red). Consistent with the d_c_ = 5 example above, this suggests that the avalanche statistics are not impacted by the activity in the dimensions defined by the high PCs. Indeed, when we reconstructed the data using PCs d_c_ through N, the power-law range dramatically dropped when d_c_ exceeded a relatively small number (Fig 1J blue). Thus, the critical subspace is strongly dependent on the first few PCs. This observation suggests a convenient quantitative definition for the dimensionality of the critical subspace - the number of PCs that can be removed (starting from PC 1) before the power-law range drops below 1.5 decades. Using this definition, we found that the critical subspace rarely had a dimensionality larger than 3 (13 at most, Fig 1K). Thus, we conclude that the critical subspace is always low dimensional and is always spanned by the first few PCs.

### Temporal coarse-graining is required to reveal critical subspace

The activity in the critical subspace manifests as large amplitude fluctuations, coordinated across many neurons. Previous studies suggest that the spatiotemporal structure of such population activity can depend on the time scale of observation (*31, 42–44*). Moreover, theory of critical phenomena suggests that temporal coarse-graining, i.e. excluding details at the small time scales, may be required to reveal universal properties of critical dynamics (*45*). Therefore, we next sought to determine how the critical subspace depends on the time scale of observation ΔT (for the results in Fig 1, ΔT = 50 ms). In many previous studies of neuronal avalanches based on spike recordings (*30, 31, 46, 47*), a common approach has been to set ΔT to the average interspike interval 〈*ISI*〉 for the entire population of recorded neurons, following the approach pioneered by (*48*). We note, however, that this approach was originally developed for LFP-peak events, not spikes. For our recordings here, the average single neuron spike rate was about 3 Hz. Thus, for a typical recording of 200 neurons, the 〈*ISI*〉 was about 1.5 ms. Obviously, 〈*ISI*〉 will be even smaller in experiments with more recorded neurons. Here we systematically investigated a range of ΔT between 1 ms and 500 ms.

First, since the critical subspace coincides with the first several PCs, we quantified how the importance of these PCs depends on ΔT. We found that the variance explained by the first 5 PCs is relatively small for small ΔT, but rises sharply around ΔT ∼ 10 ms (Fig 2A). Next, we asked how many neurons are involved in the first 5 PCs. In principle, it is possible that as few as 5 neurons are fully responsible for the first 5 PCs. We measured the fraction of neurons with strong engagement (loading > 0.1, Methods) with at least one of the first five PCs. We found that, as we increased ΔT, the fraction of neurons engaged with this low-dimensional subspace triples, reaching 30% (Fig 2B); note that 30% is around 30 to 60 neurons for our recordings. Thus, the importance of the first 5 PCs is hidden for small time scales, and emerges only after temporal coarse-graining. In our initial example with ΔT = 50 ms we saw that the population sum activity of the full population was very similar to that reconstructed from PCs 1-5 (Fig 1D,H). Next, we asked how the similarity between these two signals depends on ΔT. We found that they were not strongly correlated for small ΔT; this correlation sharply increased around ΔT ∼ 10 ms (Fig 2C).

**Fig 2.**
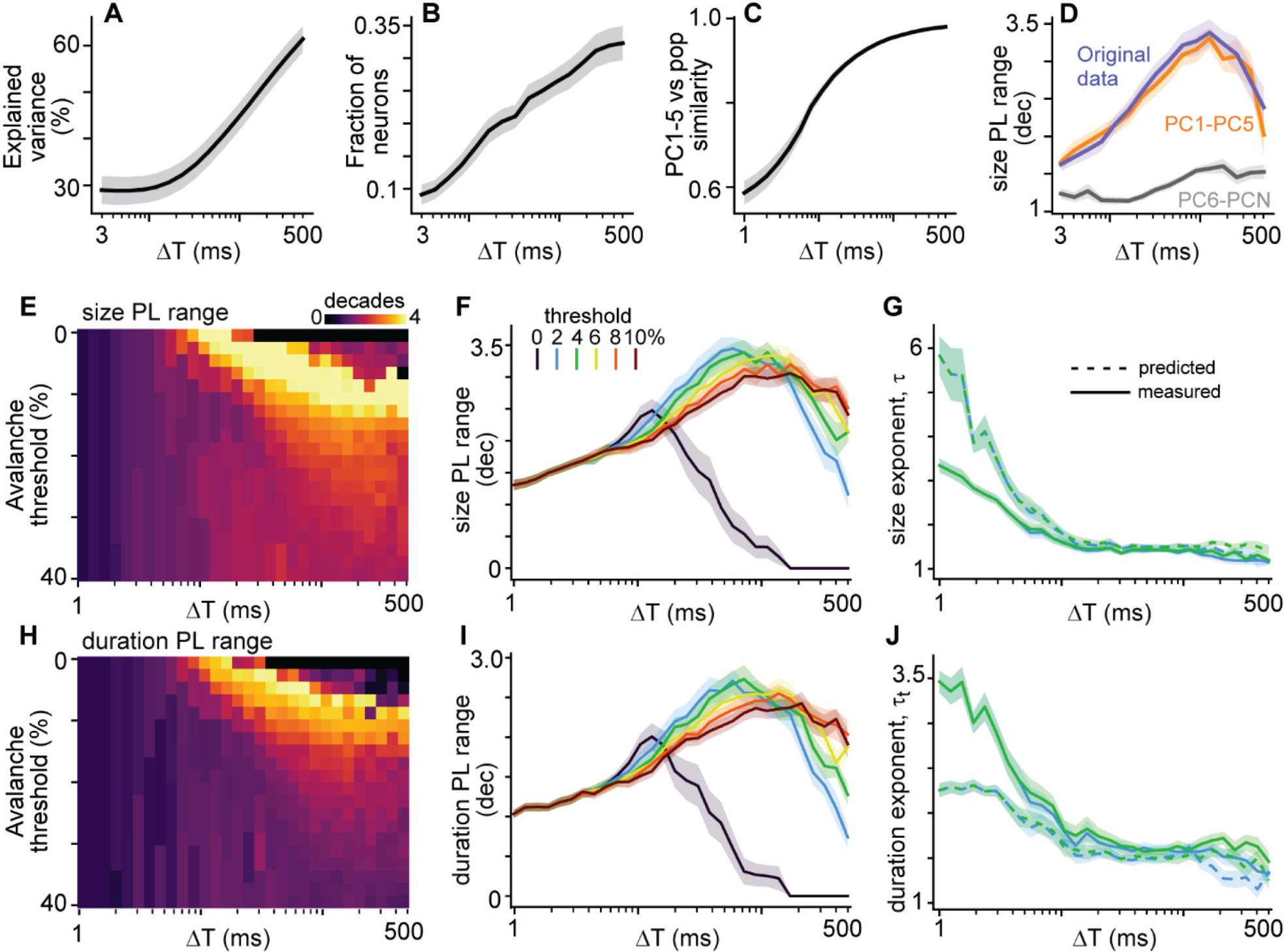
Temporal coarse-graining reveals critical subspace. **(A)** Total explained variance for the first five principal components is sensitive to temporal coarse-graining, sharply r*ISI*ng for ΔT > 10 ms, indicating the emergence of a low-dimensional subspace. **(B)**, Fraction of neurons engaged with (loading > 0.1) at least one of the first five principal components increases with ΔT. **C)** Similarity between the original summed population activity and that reconstructed using the first five principal components increases with ΔT. **(D)** Large power-law range is less than 2 decades for ΔT < 10 ms for the original data and PCs 1-5. For PCs 6-N, power-law range never exceeds 2 decades. **(E)** One example recording showing how power-law range (for population sum) depends on ΔT and the threshold used for defining avalanches. For large ΔT, the traditional approach (zero threshold) precludes finding a good power law. **(F)** Summary of how avalanche size power-law range depends on threshold and ΔT for all recordings. **(G)** The measured exponent τ (solid lines) of the size distribution power-law decreases as ΔT increases for ΔT < 10 ms, but is relatively steady for ΔT > 10 ms. Only for ΔT > 10 ms do we find agreement between the measured τ and that predicted by the crackling noise scaling law (dashed). **(H-J)** Same as panels E-G, but for avalanche duration statistics. For all panels, solid lines represent the mean across all recordings and shaded areas represent standard error.

Next, we determined how ΔT impacts evidence for criticality based on avalanche analysis. We quantified the range of good power-law fit (number of decades, Methods) for the avalanche size distribution; we interpret a larger power-law range as better evidence for criticality. For both the full population and the PC 1-5 subspace, we found that at small time scales power-law range is small (Fig 2D) and the avalanche distribution is better fit by an exponential distribution (Supplementary Fig S1). The power-law range rises around ΔT ∼ 10 ms. If we consider avalanches based on the PC 6-N subspace, power-law range is small for all ΔT (Fig 2D, grey). We note that the emergence of large power-law range at larger ΔT also depends on the threshold used for defining avalanches. Some previous studies have used a median threshold (*26, 34*), some have used the 35th percentile (*32, 33*), and many have used a zero threshold (*27, 29–31, 46, 47, 49*). Here we found that if the threshold is set to zero, the emergence of robust power-law statistics at larger ΔT will be missed (Fig 2 E,F,H,I). Finally, we examined the exponents of the power-laws and found that for ΔT < 10 ms, the exponents did not agree with the crackling noise scaling laws predicted at criticality and varied dramatically as ΔT changes (Fig 2G,J). For larger ΔT, the exponents were robust, less dependent on ΔT, and were in good agreement with crackling noise scaling laws (Fig 2G,J). Taken together, the results in Fig 2 show that evidence for criticality is robust in the critical subspace, but will be missed if spike data are not sufficiently coarse-grained in time. The critical subspace emerges for timescales above about 10 ms.

### Theory of critical dynamics confirms experimental results

Does the necessity of temporal coarse-graining to find the critical subspace agree with theory? Could the temporal coarse-graining we employ mislead us, producing apparent critical dynamics from a system that is not actually at criticality? Here we address these questions, showing that our results agree with a simple, model – the multivariate Hawkes process (*50, 51*) – but only if the model is tuned to operate close to criticality. Our model is similar to branching processes and random walks (*40, 45, 52*), providing an established theoretical case of critical dynamics, but unlike many of these classic models, the Hawkes process treats time continuously. This is helpful for our goals here of studying how the dynamics depend on the timescale ΔT. Most discrete time models can have 〈*ISI*〉 well below the duration of one time step, which precludes quantitative comparison to previous studies which often set ΔT = 〈*ISI*〉.

We set up our model with N = 10^4^ units with fixed, random connectivity. In a multivariate Hawkes process, each unit’s spikes are drawn from an inhomogeneous Poisson process. At time t, the ith unit fires with rate λ_i_(t) which depends on the recent history of spikes from other connected units,

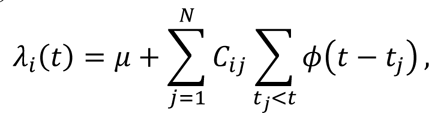

where C is the connectivity matrix (C_ij_ = 1 with probability 0.1, otherwise C_ij_ = 0). The parameter μ determines a baseline rate of noisy or spontaneous firing, independent of interactions among units. The second sum accounts for the spikes fired by the jth unit (at times t_j_) before time t and with weight that decays exponentially ϕ(*t*) = *e*^−*t*^. The magnitude of the largest eigenvalue of C, called Λ here, determines whether the system operates at criticality. For Λ well below 1, the units fire relatively asynchronously, the system is subcritical. As Λ approaches 1, firing rates and fluctuations become large and correlated and the autocorrelation diverges (Fig 3B,C); Λ = 1 at criticality.

**Fig 3.**
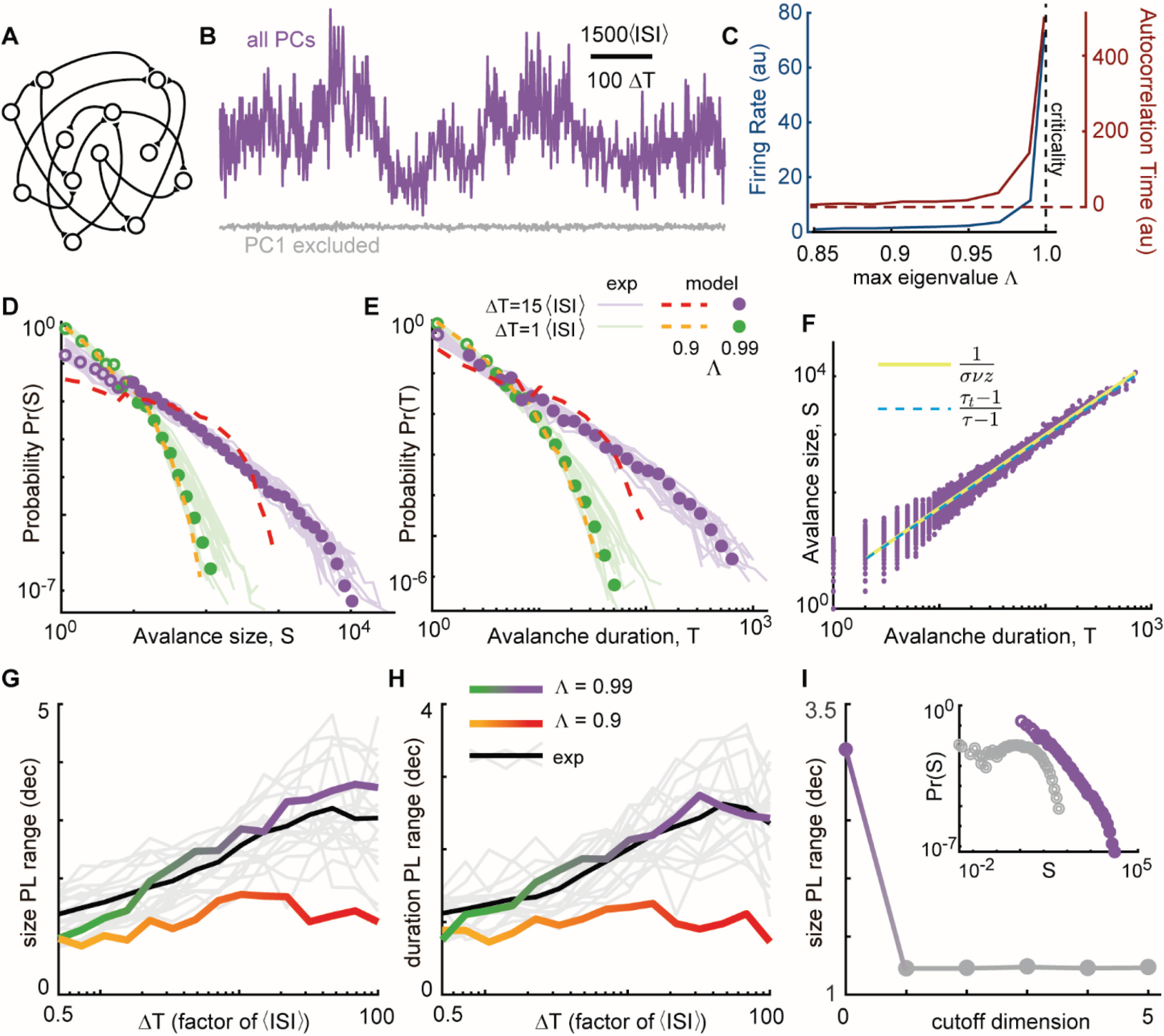
Low-dimensional model of critical dynamics agrees well with experiments. **(A)** We study a multivariate Hawkes process with random connectivity. **(B)** When the model is near criticality Λ = 0.99), the model generates population activity with large fluctuations (purple), which are abolished if the first principal component is projected out (gray). **(C)** As Λ approaches 1 (criticality), the firing rate and the timescales of fluctuations diverge. **(D)** Near criticality and with sufficient temporal coarse-graining, the model avalanche sizes were power-law distributed (ΔT = 15〈*ISI*〉, purple points), but the power-law was obscured for insufficient temporal coarse-graining (ΔT = 1〈*ISI*〉, green points). Subcritical model dynamics were not power-law distributed, with or without temporal coarse-graining (dashed lines). Experiments (light colored lines) agree well with the model. **(E)** Same as panel D, but for avalanche duration. **(F)** With ΔT = 15〈*ISI*〉, the model agrees with the crackling noise scaling law. **(G,H)** At criticality, the power-law range becomes large only for large ΔT. The subcritical model never exhibits more than 2 decades of power-law range for all ΔT. Individual experiments (gray) and mean across experiments (black) agree best with model with Λ=0.99. **(I)** The model critical subspace is 1-dimensional; power-law range collapses to less than 1.5 decades after excluding PC1. Inset shows size distributions with (purple) and without (gray) PC1.

We performed avalanche analysis on the model population activity, as we did for the experiments. We analyzed the dynamics of a subset of only 200 units, neglecting the rest, to account for subsampling effects that are certainly present in our experiments and may be important for assessing critical dynamics (*53, 54*). When the model was near criticality (Λ = 0.99), the dynamics closely matched our experimental results. Avalanche sizes and durations were power-law distributed and obeyed the crackling noise scaling law (Fig 3D-F), but only after sufficient temporal coarse-graining (approximately ΔT > 10〈*ISI*〉, Fig 3D,E,G,H, Supplementary Fig S1). When the model was subcritical (Λ = 0.9), the model dynamics were not power-law distributed for any ΔT, demonstrating that temporal coarse-graining does not produce evidence for criticality unless the system is in fact near criticality (Fig 3D,E,G,H, Supplementary Fig S1). When the noise parameter μ was increased, the importance of temporal coarse-graining increased (Supplementary Fig S1). We note that when we coarse-grained our experimental data using a ΔT set to a specified factor of 〈*ISI*〉 (instead of a specified number of milliseconds), the results of different experiments were more consistent with each other and in good quantitative agreement with the model (light colored lines in Fig 3D,E,G,H).

Like the experiments, critical dynamics in the model are confined to a low-dimensional subspace. In fact, the model is essentially 1-dimensional. Removing the first principal component fully removes the collective fluctuations in firing rates (Fig 3B). Moreover, removing the first PC completely abolishes the power-law statistics of avalanches (Fig 3I). In Supplementary Fig S2, we show that the model can generate higher dimensional dynamics if we implement spatially-dependent near-neighbor connectivity, similar to critical phenomena in other systems with spatial translational invariance (*55*). However, the neurons recorded from one high-density shank electrode like in our experiments are all within a few hundred micrometers from each other and, thus, are expected to have more spatially independent connectivity. Thus, we conclude that our experimental measurements are well-explained by theory of low-dimensional critical dynamics.

### Desynchronized subspace coexists with the critical subspace

As discussed in the introduction, the standard point of view is that criticality is not compatible with desynchronized dynamical regimes; it is thought that the cortex must “choose” between criticality and a desynchronized regime. However, the fact that the critical subspace is low-dimensional raises an interesting question that challenges the standard view. Could there be a desynchronized dynamical regime that coexists with the critical subspace, in the same neural circuit, but in a different subspace?

As demonstrated in Figs 1C,J, and K, the number of dimensions needed to explain a substantial fraction of the total variance is much greater than the dimension of the critical subspace. This suggests that the additional dimensions beyond the critical subspace are “important”, in the sense that they are needed to explain much of the measured variance. What type of dynamics is contained in these additional dimensions? The basic mathematical facts of PCA require that these dimensions outside the critical subspace must have smaller fluctuations, because the critical subspace spans the first few PCs. But PCA does not tell us whether the dynamics in these dimensions has the characteristic features of a desynchronized regime: weak correlations among units, short autocorrelation timescales, and Gaussian distributed population activity. To answer these questions we reconstructed our measured spike count dynamics using a small number of PCs (d_c_+1 through 2d_c_) just beyond the cutoff dimension d_c_ of the critical subspace, as defined in Fig 1J,K. Fig 4A shows an example recording where each subspace is 6-dimensional (i.e. d_c_ = 6).

**Fig. 4.**
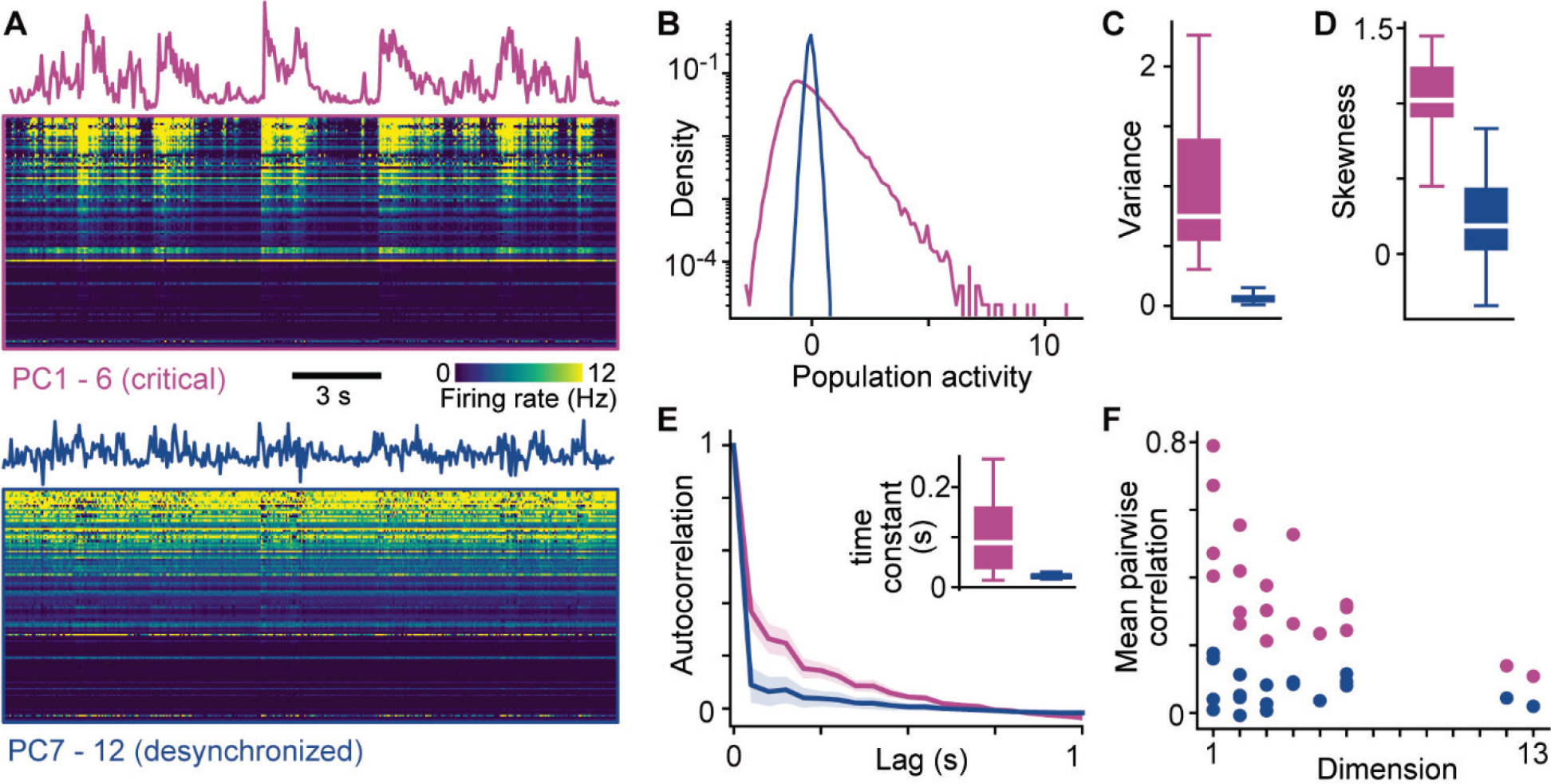
Desynchronized subspace coexists with critical subspace. **(A)** Population summed activity and raster reconstructed from the critical subspace (purple, d_c_=6 in this example) and the desynchronized subspace (blue). **(B - D)** The distribution of population summed activity is broad, heavy-tailed, and skewed for the critical subspace, but narrow and near gaussian for the desynchronized subspace. Box-plots summarize all experiments (box spans quartiles, white line is median). **(E)** For the desynchronized subspace, the autocorrelation function drops sharply indicating short-range temporal correlations compared to the critical subspace. Inset: time constants from an exponential fit for all experiments. **(F)** Mean pairwise correlations among reconstructed single unit activities are closer to zero for the desynchronized subspace.

First, we summed the reconstructed dynamics across units to obtain a single population summed activity for each subspace (Fig 4A). As expected, this population activity fluctuated with much larger amplitude (larger variance, Fig 4B,C) in the critical subspace compared to the desynchronized subspace. In addition, we found that the distribution of population activity in the desynchronized subspace was close to Gaussian (low skewness), while the critical subspace had heavy tailed, skewed distributions (Fig 4D). Another defining feature of desynchronized regimes is a lack of long-range temporal correlations. We compared autocorrelation functions for the population activity in the two subspaces and found that the critical subspace had much longer temporal correlations (Fig 4E). Finally, we assessed pairwise correlations among units, using the reconstructed spike count rasters (Fig 4A). Here, some caution is needed; the dimensionality of the subspace impacts the basic nature of pairwise correlations for such reconstructed unit activity. Nonetheless, defining the dimensionality of the two subspaces to be equal allows for meaningful relative comparisons and we found that the critical subspace always had much higher pairwise correlations than the desynchronized subspace. Thus we conclude that the dynamics in this subspace with the greatest variance, after removing the critical subspace, manifested all the defining features of desynchronized regimes: weak pairwise correlations, short-range temporal correlations, and Gaussian distributed population activity (*9, 56*).

## Discussion

We have shown that population spiking activity in awake mouse motor cortex can be partitioned into different subspaces, each containing fundamentally different kinds of coordinated dynamics. The most prominent subspace – defined by the first few principal components – is home to critical dynamics with long-range temporal correlations, heavy-tailed distributions of activity and multifaceted agreement with scaling laws predicted at criticality. The next most prominent dimensions contain desynchronized dynamics with weak spatiotemporal correlations and Gaussian distributed population activity.

These observations open several interesting new questions and offer answers to several long-standing questions about the fundamental operating regime of cortical neuronal networks. One such long-standing question asks whether cortical circuits operate near criticality or not in awake cortex. Many previous experiments with strong agreement with criticality theory were based on techniques that measure collective brain signals, like local field potential (LFP) (*30, 48, 57–60*), wide field imaging (*25, 61*), and human brain imaging (*57, 62*). These collective signals represent an aggregate of the underlying spike activity of many individual neurons. However, when spikes recorded in awake animals have been analyzed directly, results have been less clear - some studies report support for criticality (*31, 33–38, 49, 63*) while others do not (*27–32*). Considering that spikes are the fundamental information carriers underlying brain function, the equivocal support for criticality at the level of spike measurements has created skepticism and confusion surrounding the hypothesis (*29, 30*). Why is evidence for criticality clear in collective signals, but unclear in spike data? As we showed here, one important factor explaining discrepancies among previous spike recordings is that the existence of the critical subspace is clear only at larger timescales, beyond about 10 〈*ISI*〉. Our results suggest that the studies based on spikes with negative reports about criticality missed the evidence for criticality because they did not sufficiently coarse-grain in time (Supplementary Table 1). What about collective signals like LFP? One possibility is that collective signals, by their nature, may carry some degree of overlapping signals (LFP at two nearby locations can reflect signals from the same source neurons), which results in spurious correlations that could be mistaken for criticality (*64*). Considering our results here, another possibility is that the coordinated activity in the critical subspace is apparent in measurements of collective signals like LFP, but the desynchronized subspace is hidden from collective signals because it is weakly coordinated. Thus, measuring collective signals may effectively filter out the desynchronized subspace, leaving only the critical subspace signals.

Several recent studies have proposed that critical phenomena in neural systems might be fundamentally high dimensional (*65–69*). Do these studies contradict our work here? Several of these previous studies focused on edge-of-chaos (EOC) criticality, which, unlike our results here, is thought to occur without large fluctuations at the population level (*66, 68, 69*). In other words, EOC criticality has no avalanches. In this basic sense, our experimental results are inconsistent with EOC criticality – we observe prominent large amplitude fluctuations and power-law distributed avalanches. In other studies (*55, 65–67*), criticality was hypothesized to be high-dimensional based on data analysis and concepts adapted from traditional renormalization group in systems with spatial translational invariance (similar to the *ISI*ng model, for example). Our experimental results do not directly confirm or deny this hypothesis, but our model shows a clear case of low-dimensional criticality; thus, criticality in neural systems is not required to be high dimensional. We further clarify this point in Supplementary Fig S2; we show how a transition from low dimensional to high dimensional criticality can result from tuning the model connectivity from global to local and introducing approximate spatial translational invariance. These theoretical considerations suggest that we observed a low dimensional critical subspace in our experiments, because we measured neurons from a local patch of cortex (approximately one cortical column) using a single shank Neuropixels probe. Within the population of recorded neurons connectivity is approximately global. Thus, we might expect to find a higher dimensional critical subspace based on more spatially wide-spread recordings, as found in recent fMRI measurements, for example (*55*).

As discussed in the introduction, another controversy that our results may help resolve is the question of cortical state: Are cortical population dynamics weakly or strongly correlated? Our results show that this question may be a false dichotomy; we show that weakly correlated dynamics can coexist with relatively strongly dynamics by separating them into different subspaces. However, this observation raises important follow up questions. Do these two different subspaces perform different brain functions? Can the different subspaces really be used by the brain in a way that confers the different functional advantages of weakly versus strongly correlated dynamics? For example, desynchronized dynamics are thought to be beneficial for low-noise discrimination and representation of sensory input. Whereas criticality is thought to be beneficial for multi-scale information transmission, to name one example. Do previous studies support the possibility that the critical subspace and the desynchronized subspace might execute different functions? Stringer et al (2019) showed that visual input is encoded in one subspace with relatively small fluctuations while some body movements (e.g. running and whisking) are encoded in a different subspace with larger fluctuations (first few PCs). Moreover, Jones et al (2023) reanalyzed the data from Stringer et al (2019) and found that subsets of neurons with the strongest correlations to body movements also exhibited power-law fluctuations similar to the avalanches we observed in our critical subspace here. Taken together, these results suggest that sensory input might be encoded in the desynchronized subspace we found here, while behavior may be encoded in the critical subspace. Another possible use for the critical subspace might be exchange of signals among cortical areas (*70, 71*). These studies used ongoing activity in visual cortex (i.e. they studied the stimulus independent “noise” fluctuations, consistent with our critical subspace). Further studies will be needed to carefully test these interesting possible functional implications.

## Materials and Methods

## Animals

All procedures followed the Guide for the Care and Use of Laboratory Animals of the National Institutes of Health and were approved by University of Arkansas Institutional Animal Care and Use Committee (protocol 21022). We studied adult male C57BL6/6J mice (Jackson Labs). After acclimatization to handling, a small aluminum plate (0.5 g), was attached to the skull with dental cement. Then, mice were trained for head fixation for 20 sessions, gradually increasing in duration. At the time of recordings, the mice weighed ≈ 28 g and were 21-23 weeks old. 1-2 days before the first recording for each mouse, a craniotomy (2 mm diameter) was performed over right motor cortex (Anterior-Posterior = 0 mm, Medial-Lateral = 1 mm). Each recording day began with a brief period of isoflurane anesthesia to expose the craniotomy and head fix the mouse. The mice were free to run, sit, groom, and walk for the entire duration (45 minutes) of each recording. During recordings, after inserting the electrode array, the craniotomy was covered with gel-foam pieces soaked in sterile phosphate buffer solution.

## Electrophysiology

The extracellular voltage was recorded using Neuropixels probes (NP version 1.0, IMEC) consisting of an electrode shank (width: 70 μm, length: 10 mm, thickness: 100 µm) of 960 total sites laid out in a checkerboard pattern with contacts (18 µm site-to-site separation), enabling up to 384 recording channels. On the recording day, following head fixation, the Neuropixels probe was inserted to a tip-depth of approximately 1.2 mm, ensuring that the active recording sites spanned all cortical layers. An Ag/AgCl pellet was used as ground, placed in the saline-soaked gel foam covering the craniotomy. The ground pellet wire was soldered to the Neuropixel midway along the ribbon cable. Electrophysiological data were collected (30 kHz) using SpikeGLX software. Spike sorting was performed using Kilosort 3.0 (https://github.com/MouseLand/Kilosort) and then manually curated using phy (https://github.com/cortex-lab/phy) (*72*).

## Data analysis

*PCA* - We performed principal component analysis (PCA) in Python using the function ‘decomposition.PCA’ from package ‘sklearn’. Let Z be a spike count matrix with M rows (number of time bins) and N columns (number of neurons). Then, PCA generates V which contains the principal components, i.e. the eigenvectors of the covariance matrix of Z. V has N rows and N columns (one column for each eigenvector, i.e. one column for each principal component). We calculated % variance explained by a set of PCs as the sum of their corresponding eigenvalues of V (reported in Fig 1C and Fig 2A). To reconstruct data based on a subset of PCs (e.g. PCs 1-K), we first define the M × K projection matrix B by B=Z*V^^^*, where *V^^^*is N × K, including the first K columns of V. Then the reconstructed activity is 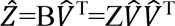.

*Avalanche analysis* - The first step in avalanche analysis was to create a spike count matrix Z, where Z_tj_ is the number of spikes fired by unit j during time bin t. Next, the population sum spike count time series X was created by summing spike counts over all neurons at each time bin, *X*_*t*_ = ∑_*j*_ *Z*_*tj*_. The threshold ϴ used for avalanche detection was defined as some percentile of X. By definition, an avalanche begins when X exceeds the threshold and ends when X returns below threshold. The size S of an avalanche is defined 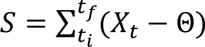, where the start and end times of the avalanche are t_i_ and t_f_, respectively. Avalanche duration is defined as T = t_f_ - t_i_.

In order to assess whether avalanche sizes and durations were distributed according to a power-law and to obtain power-law exponents and ranges, we built on previously developed maximum likelihood methods (*34, 58, 73, 74*). In brief, the fitting algorithm identifies the best fit truncated power-law that meets a pre-defined goodness-of-fit criterion. There are three fitting parameters: the minimum avalanche size x_m_, the maximum avalanche size x_M_, and the power-law exponent τ. The following steps summarize the algorithm. First, events with size/duration less than x_m_ and larger than x_M_ were excluded. Second, the maximum likelihood power-law exponent was calculated. Third, we assessed the goodness-of-fit. We repeated these four steps for all the possible pairs of x_m_ and x_M_ values, in the end, identifying the largest power-law range that passed the goodness-of-fit criterion. We define power-law range as the number of decades of power-law scaling log_10_(*x*_*M*_/*x*_*m*_). We note that this algorithm is independent of any choice of bins used to create the PDF plots in the paper.

The primary improvement we made compared to our most recently published methods (*34*) was to make our goodness-of-fit criterion less sensitive to sample size (number of avalanches) and more computationally efficient. For a given x_m_, x_M_, and τ, goodness-of-fit was quantified as follows. First, we created a cumulative distribution function (CDF) of the real data (excluding samples below x_m_ and x_M_). Second, we define a theoretical CDF for a truncated power-law with the same range and exponent. Third, we define a region delimited by upper and lower bounds defined as the theoretical CDF +0.03 and -0.03, respectively. Fourth, we resample the real CDF at 10 logarithmically spaced values per decade. Fifth, we calculated the fraction F of resampled points in the CDF of the real data that fell within ±0.03 bounds of the theoretical CDF. F is our goodness-of-fit measure. F=1 means that the entire range of the real data varies less than 3% from a perfect power-law. We sought the fit with largest power-law range that meets the goodness-of-fit criterion F ≥ 0.8.

*Decorrelated subspace* – The analysis presented in Fig 4, began with making a spike count matrix with time bin size ΔT = 50 ms. Next, the activity was reconstructed using different dimensions as described above. The third step was to perform a band pass filter including 0.1 – 100 Hz on the reconstructed activity time series for each unit. Finally, the activity distributions, activity variance and skewness, autocorrelation functions, and pairwise correlations were analyzed. The time constant for the autocorrelation functions was the time constant of a best-fit exponential function.

## Computational model

The numerical implementation of the multivariate Hawkes process described in Fig 3 and corresponding text was done according to ref (*75*).

## Acknowledgments

**Funding:**

National Institutes of Health grant R15NS116742 (AJF, VKN, SHG, WLS)

National Science Foundation grant R01AB123456 (AJF, SHG, WLS)

Arkansas Biosciences Institute (SHG, WLS)

## Author contributions

Conceptualization: AJF, JSS, WLS

Experimental recordings: AJF, VKN, SHG, WLS

Data analysis: AJF, JSS, WLS

Computational modeling: JSS

Figure creation: AJF, JSS, WLS

Writing: WLS, AJF, JSS

## Data and materials availability

Spike data and analysis code are publicly available without restriction on Figshare at doi: *TBD upon paper acceptance*. All data are available in the main text or the supplementary materials.

## Supplementary Materials

Supplementary Materials include Supplementary Text, two Supplementary Figures, and one Supplementary Table.

## Supplementary Materials for

### Supplementary Text

#### Temporal coarse-graining reveals power-laws

In the main text, we presented power-law distributed avalanches after sufficient temporal coarse-graining and claimed that avalanche distributions without coarse-graining were not power laws. Here directly support this claim, quantitatively showing that when ΔT is too small, the avalanche distributions are better fit by an exponential distribution than a power-law. We compared the quality of the maximum likelihood exponential fit (*P*_*exp*_(S) ∘ e^−S/a^, fitting parameter a) to that of the maximum likelihood power-law fit (*P*__*pl*__(S) ∘ S^−τ^, fitting parameter τ) using the Akaike information criterion (AIC) (Fontenele et al., 2019; Hirotugo, 1974).

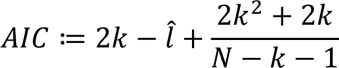

where k = 1 is the number of fitting parameters, N is the number of samples, and *l*^^^ is the log-likelihood at its maximum. Unlike in the power-law fitting algorithm used in the main text, here we included all observed avalanches in the fit, without imposing an upper or lower cutoff. Smaller AIC means a better fit, so Δ ≔ *AIC_pl_* – *AIC_exp_* is greater than zero when the exponential fit is better than the power-law fit, and less than zero when the reverse is true. As shown in Fig S1, the exponential was a better fit for all experiments except 1 when ΔT=〈*ISI*〉 and the power-law was a better fit for all experiments when ΔT=30〈*ISI*〉.

#### Dimensionality of critical dynamics depends on connectivity of units

In the main text Discussion, we mentioned that some previous studies have hypothesized that critical dynamics in neural systems is high dimensional, unlike the low-dimensional critical subspace we presented. Here we show using our computational model, that the dimensionality of critical dynamics depends strongly on the nature of connectivity among neurons in the model. The main results in Fig 3 are based on global random connectivity; each pair of neurons is connected with probability p = 0.1, independent of the distance between the two neurons. In this case, the critical dynamics are 1-dimensional as we showed in Fig 3. However, if we consider neurons on a square lattice (50 x 50) with local connectivity (connection probability is strongly dependent on distance between neurons), then we find that the critical dynamics of the model become higher dimensional (Fig S2). The spatial reach of local connectivity is inversely related to the dimensionality of the critical subspace (computed in the same way as in Fig 1 of the main text).

**Fig. S1.**
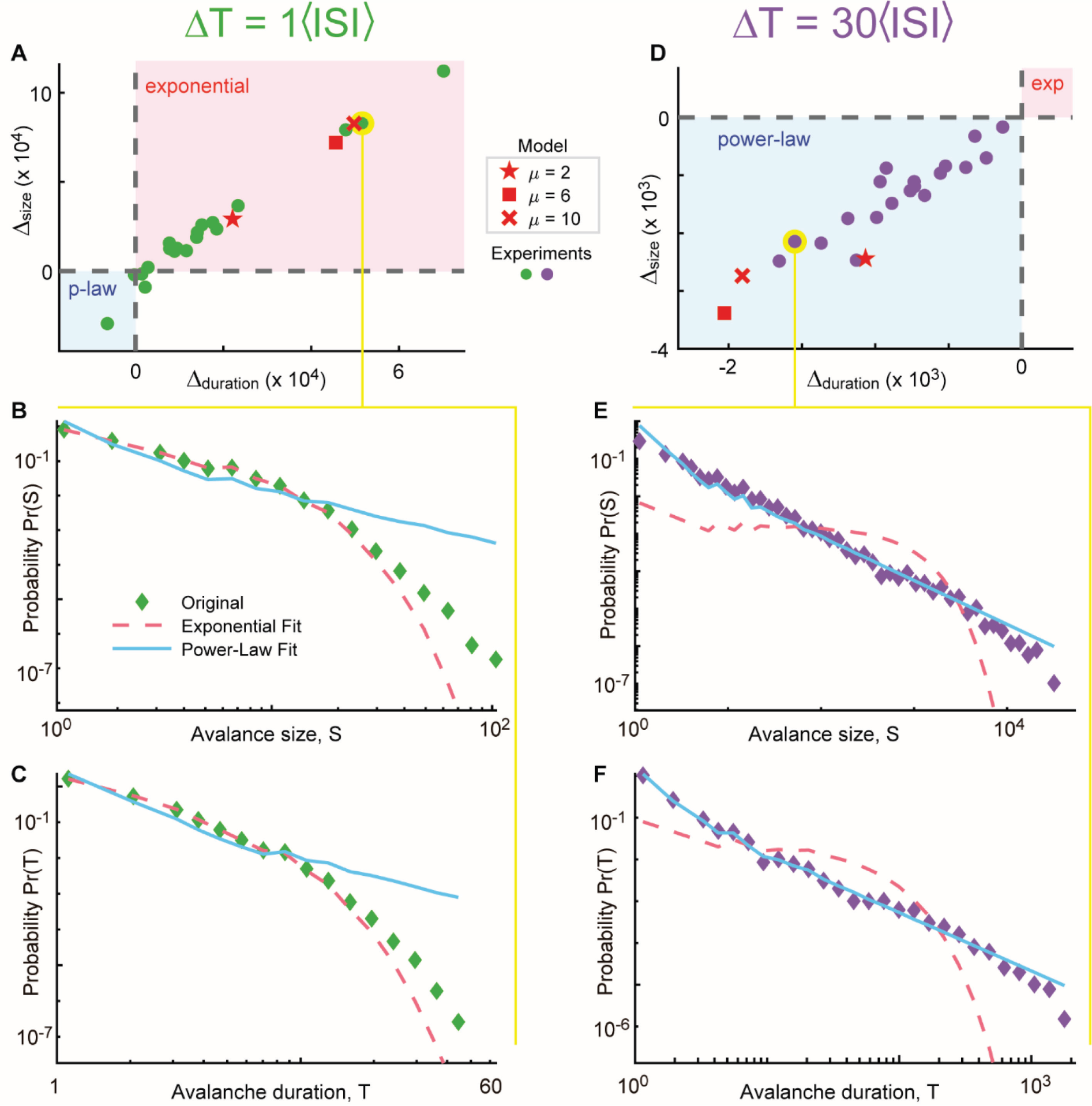
Without temporal coarse graining, avalanches are exponentially distributed. (**A**) Each point represents one experiment (green) or one run from the model (red) analyzed using ΔT=〈*ISI*〉. The vertical position of each point represents the comparison Δ ≔ *AIC*__*pl*__ − *AIC*_*exp*_for the avalanche size distribution; the horizontal position represents the same comparison, bur for the duration distribution. A point that falls in the pink shaded area is best fit by an exponential distribution for both size and duration. Note that, for the model, increasing uncorrelated noise (μ) results in better exponential fits, suggesting that uncorrelated firing can be one reason that temporal coarse graining is needed. (**B,C**) The size and duration distributions for the example point highlighted in yellow in panel A. Clearly, the exponential fit (dashed) is better than the best power-law fit (blue line). (**D-F**) Same as panels A-c, but ΔT= 30〈*ISI*〉, resulting in all experiments being better fit by a power-law for both size and duration distributions (blue shaded region). Note: the blue lines are perfect power laws. The slight deviations from a straight line on log-log axes are due to logarithmic binning to create the PDF plots.

**Fig. S2.**
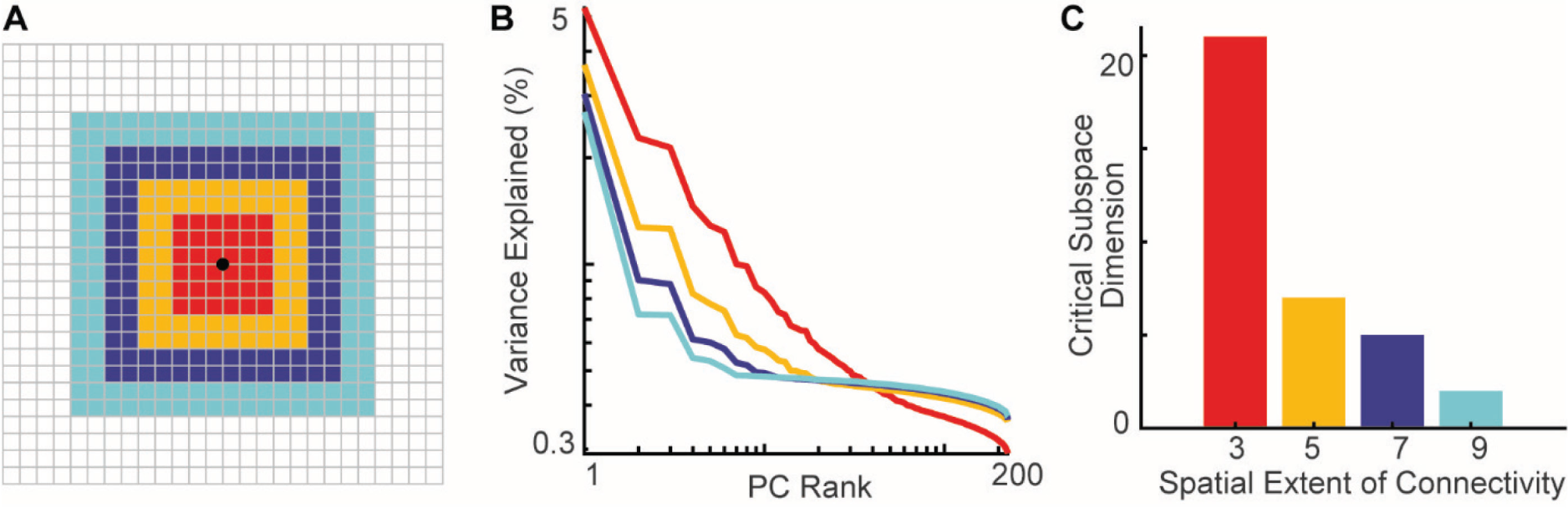
Spatial translational invariance and local connectivity results in higher dimensional critical subspace. (**A**) We simulated a multivariate Hawkes Process on a 50 x 50 lattice, with outgoing connections from each site to the sites in an L x L square centered on it. All nonzero connections are excitatory and of equal strength. As in the main text, we scaled the connectivity matrix to have largest eigenvalue Λ = 0.99, po*ISI*ng the network close to criticality, and set the baseline spiking rate to μ = 6. For the avalanche analysis and PCA, we took a 15 x 15 localized subsample and chose a time bin of length ΔT = 30〈*ISI*〉. Increasing the spatial extent L of the connectivity decreased both the dimension of the critical subspace (**C**) and the number of PCs over which the PC eigenspectrum decays as an approximate power-law (**B**).

**Table S1.**
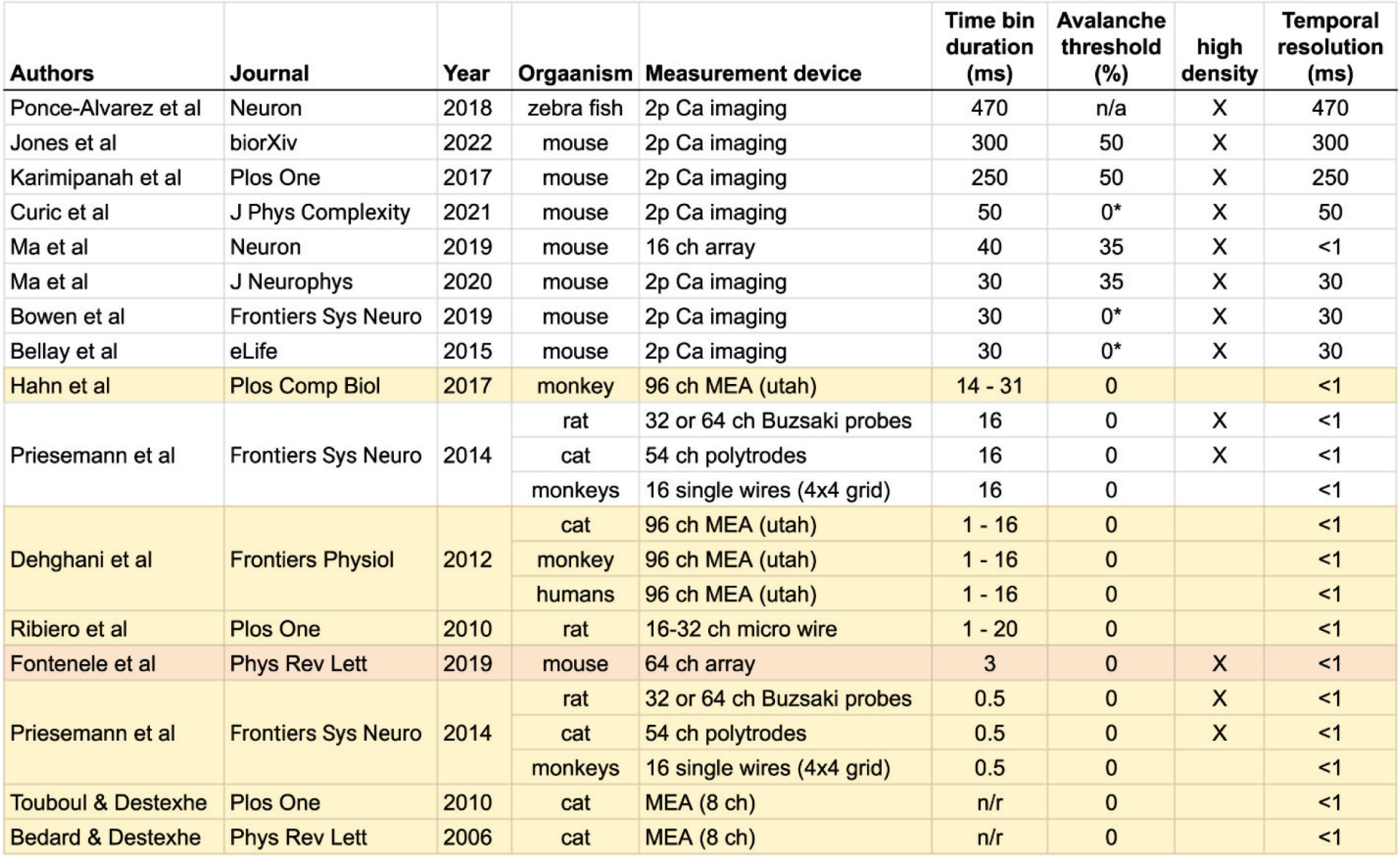
Insufficient temporal coarse-graining explains previous discrepancies. We summarize all previous experiments that performed avalanche analysis based on spikes recorded from awake animals. Experiments that reported poor evidence for power-law distributed avalanches are shaded yellow. The rows of the table are ordered from top to bottom in order of largest to smallest amount of temporal coarse graining. These results suggest that previous negative reports are due to insufficient temporal coarse-graining.

